# Functionally distinct high and low theta oscillations in the human hippocampus

**DOI:** 10.1101/498055

**Authors:** Abhinav Goyal, Jonathan Miller, Salman E. Qasim, Andrew J. Watrous, Joel M. Stein, Cory S. Inman, Robert E. Gross, Jon T. Willie, Bradley Lega, Jui-Jui Lin, Ashwini Sharan, Chengyuan Wu, Michael R. Sperling, Sameer A. Sheth, Guy M. McKhann, Elliot H. Smith, Catherine Schevon, Joshua Jacobs

**Affiliations:** Mayo Clinic College of Medicine and Science, Mayo Clinic, Rochester MN, 55905; Department of Biomedical Engineering, Columbia University, New York, NY 10027; Department of Neurology, University of Texas, Austin; Department of Radiology, University of Pennsylvania, Philadelphia, PA 19104; Department of Neurological Surgery, Emory University, Atlanta, GA 30322; Department of Neurological Surgery, University of Texas Southwestern, Dallas, TX 75390; Department of Neurological Surgery, Thomas Jefferson University, Philadelphia, PA 19107; Department of Neurology, Thomas Jefferson University, Philadelphia, PA 19107; Department of Neurological Surgery, Baylor College of Medicine, Houston, TX, 77030; Department of Neurosurgery, Columbia University Medical Center, New York, NY 10032; Department of Neurosurgery, University of Utah, Utah; Department of Neurology, Columbia University Medical Center, New York, NY 10032; Jefferson Comprehensive Epilepsy Center, Jefferson University Hospitals, Philadelphia, PA, USA

## Abstract

Based on rodent models, researchers have theorized that the hippocampus supports episodic memory and navigation via the theta oscillation, a ~4–10-Hz rhythm that coordinates brain-wide neural activity. However, recordings from humans have indicated that hippocampal theta oscillations are lower in frequency and less prevalent than in rodents, suggesting interspecies differences in theta’s function. To characterize human hippocampal theta, we examined the properties of theta oscillations throughout the anterior–posterior length of the hippocampus as neurosurgical subjects performed a virtual spatial navigation task. During virtual movement, we observed hippocampal oscillations at multiple frequencies from 2 to 14 Hz. The posterior hippocampus prominently displayed oscillations at ~8-Hz and the precise frequency of these oscillations correlated with the speed of movement, implicating these signals in spatial navigation. We also observed slower ~3-Hz oscillations, but these signals were more prevalent in the anterior hippocampus and their frequency did not vary with movement speed. Our results converge with recent findings to suggest an updated view of human hippocampal electrophysiology. Rather than one hippocampal theta oscillation with a single general role, high-and low-theta oscillations, respectively, may reflect spatial and non-spatial cognitive processes.

## Introduction

The theta oscillation is a large-scale network rhythm that appears at ~4–10 Hz in rodents and is hypothesized to play a fundamental role in mammalian spatial navigation and memory (Kahana et al., 2001; Buzsáki, 2005). However, in humans, there is mixed evidence regarding the relevance and properties of hippocampal theta. Some studies in humans show hippocampal oscillations at 1–5 Hz that have similar functional properties as the theta oscillations seen in rodents (e.g., Arnolds et al., 1980; Jacobs et al., 2007; Vass et al., 2016; Watrous et al., 2011; Watrous, Lee, et al., 2013; Jacobs, 2014). There is also evidence that human movement-related hippocampal theta oscillations vary substantially in frequency according to whether a subject is in a physical or virtual environment (Aghajan et al., 2016; Bohbot et al., 2017; Yassa, 2018). Together, these studies have been interpreted to suggest that the human hippocampus does show a signal analogous to theta oscillations observed in rodents but that this oscillation is more variable and slower in frequency (Jacobs, 2014). These apparent discrepancies in the frequency of theta between species and behaviors shed doubt on the notion that theta exists as a single general oscillatory phenomenon that coordinates brain-wide neural activity consistently across species and tasks.

Our study aimed to resolve these discrepancies by characterizing the properties of human hippocampal oscillations in spatial cognition. We analyzed intracranial electroencephalographic (iEEG) recordings from the hippocampi of fourteen neurosurgical subjects performing a virtual-reality (VR) spatial navigation task, in which subjects were asked to remember the location of an object as they were moved along a linear track (Fig. 1). A distinct feature of our experimental design compared to previous work is that our task randomly varied the subjects’ movement speed along the virtual track. This design encouraged subjects to continually attend to their spatial location throughout movement, because a non-spatial strategy based on remembering the time delay to each object would not be viable due to the speed changes within trials. We hypothesized that this feature of our task would more clearly elicit human hippocampal oscillations specifically related to navigation. Additionally, we recorded signals at various positions along the anterior–posterior axis of the hippocampus, including sites located considerably more posterior than those seen in previous work of this type, which allowed us to probe the anatomical organization of these oscillations.

**Figure 1:**
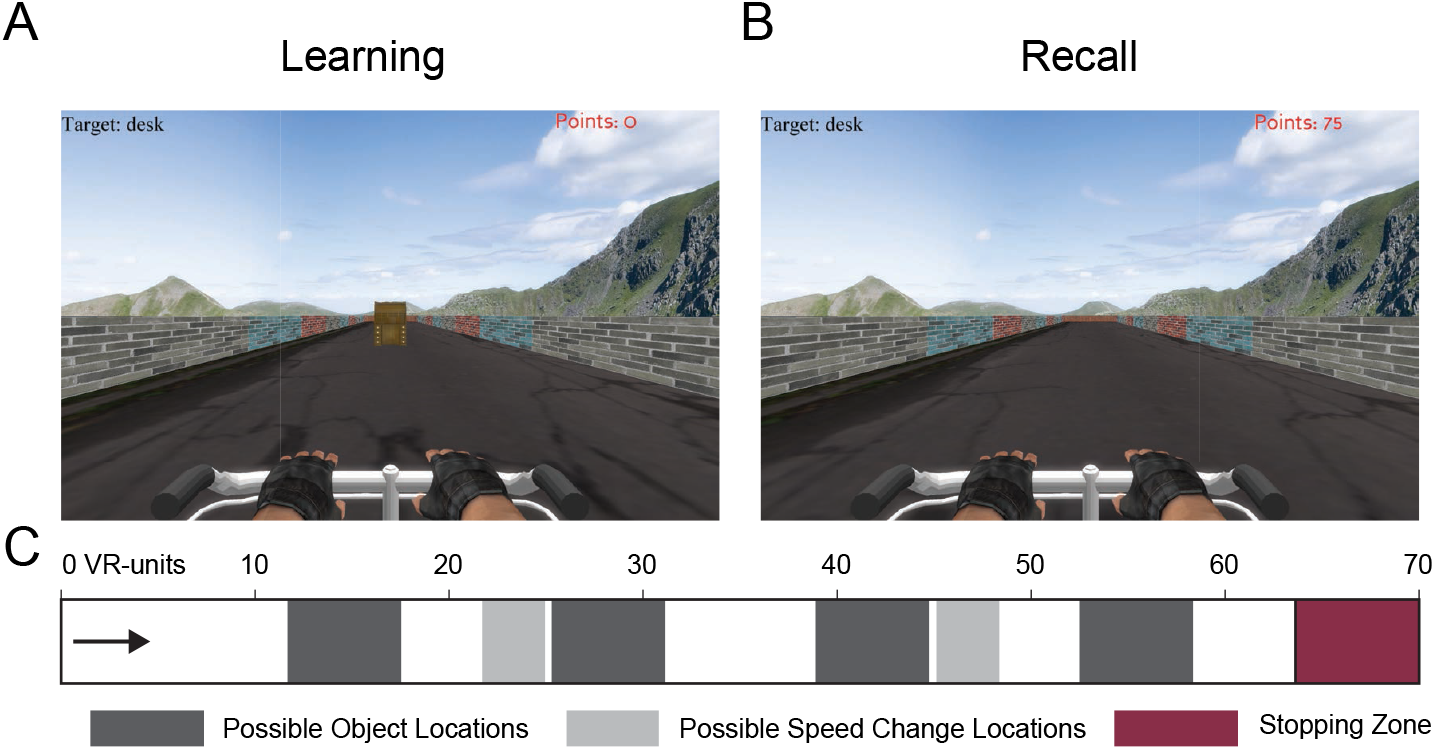
Spatial memory task. **A.** Task screen image during a learning trial, where the object is visible as the subject travels down the track. **B.** Task image during a recall trial, in which the object is invisible and the subject must recall the object location. **C.** Task schematic, showing possible object and speed change locations.

Given the anatomical differences in the hippocampus between rodents and humans (Colombo et al., 1988; Amaral & Witter, 1989; Moser & Moser, 1998; Strange et al., 2014), we hypothesized in particular that understanding the spatial organization of human theta could help explain the apparent interspecies differences that have been reported previously. Therefore, we analyzed the spectral and functional features of human hippocampal oscillations and tested their consistency along the length of the hippocampus. In contrast to earlier work that generally emphasized a single theta oscillation for a given behavior, we instead found that the hippocampus showed multiple oscillations at distinct frequencies (often at ~3 Hz and ~8 Hz), even in a single subject. Further, ~8-Hz oscillations in the posterior (but not anterior) hippocampus often correlated with spatial processing. By demonstrating multiple patterns of hippocampal oscillations with different anatomical and functional properties, our findings suggest that human hippocampal oscillations at different frequencies are generated by separate anatomical networks to support distinct functions.

## Results

Fourteen neurosurgical subjects (8 males and 6 females, age range 23–49) performed our virtual-reality (VR) spatial memory task as we recorded neural activity from iEEG electrodes implanted in their hippocampi. The task (Qasim et al., 2019) required that subjects press a button to indicate when they were located at the position of a specified hidden object as they were moved at a randomly varying speed in one direction along a linear track (Fig. 1). Overall, subjects performed the task well, responding accurately on 84% of trials (defined as having an error distance less than 11.5 VR units; see *Methods*). We performed spectral analyses of the iEEG signals during movement phases of the task for all hippocampal recording sites and used an oscillation-detection procedure known as MODAL (Watrous et al., 2018) to identify prominent narrowband oscillations (see *Methods*). Overall, we observed hippocampal narrowband oscillations at frequencies in the range of 2 to 14 Hz (Fig. 2A), consistent with earlier findings (Ekstrom et al., 2005; Jacobs et al., 2007; Watrous et al., 2011; Lega et al., 2011; Bush et al., 2017), with oscillations being most prevalent at ~3 Hz and ~8 Hz. For convenience, we label the oscillations in our dataset as low theta (2–4 Hz) and high theta (4–14 Hz) frequency bands, although we acknowledge that some other studies have used the terms “delta” and “alpha” to refer to parts of these bands.

**Figure 2:**
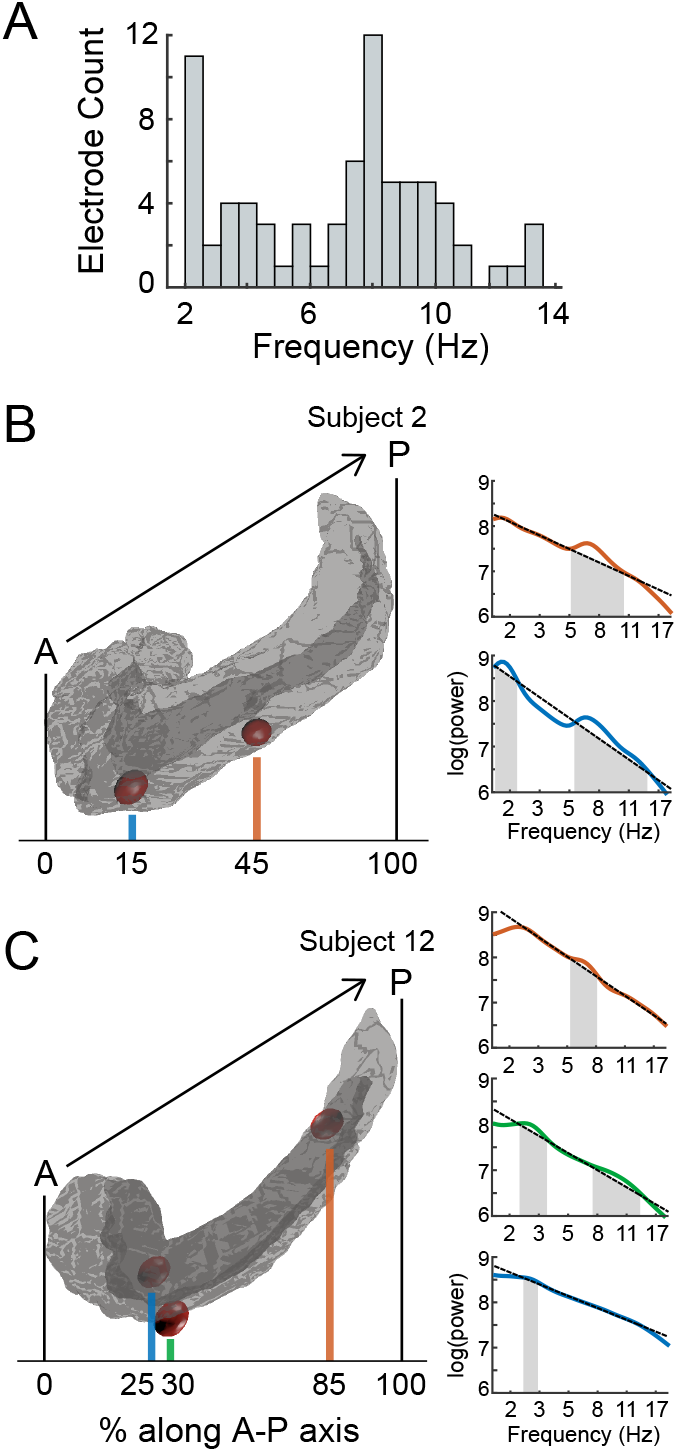
Power spectra of electrodes at different positions along the anterior–posterior axis of the hippocampus. **A.** The distribution of detected oscillations across all hippocampal electrodes in our dataset. **B.** Rendering of Subject 2’s left hippocampus (left) and the power spectra (right) for electrodes implanted in this area. Shading in the power spectrum indicates detected narrowband oscillations. **C.** Rendering of Subject 12’s left hippocampus and power spectra for the implanted electrodes.

### Anatomical organization of hippocampal high-and low-theta oscillations

We next examined how the characteristics of the oscillations we identified varied with their location along the hippocampal anterior-posterior axis. Many previous electrophysiological studies in rodents generally focused on hippocampal oscillations in the dorsal area (analogous to the posterior hippocampus of humans; Colombo et al., 1988; Amaral & Witter, 1989; Moser & Moser, 1998; Strange et al., 2014) or those that are consistent across the length of the hippocampus (Lubenov & Siapas, 2009). However, a different line of work in humans (Maguire et al., 1997; Greicius et al., 2002; Kumaran et al., 2009; Poppenk et al., 2013; Lin et al., 2018) and animals (Moser & Moser, 1998; Royer et al., 2010; Fanselow & Dong, 2010; Hinman et al., 2011) showed functional variations along the length of the hippocampus. This suggested to us that oscillations at different anterior–posterior positions could have distinct spectral and functional properties.

We measured the anterior–posterior location of each hippocampal electrode in a subject-specific manner, defined as the relative distance between the anterior and posterior extent of the hippocampus (see *Methods*). In this scheme, positions 0% and 100% correspond to electrodes at the anterior and posterior tips of the hippocampus, respectively. As seen in Figure 2B&C, within individual subjects, we observed narrowband oscillations at various frequencies. Individual electrodes displayed oscillations at either one or two distinct frequency ranges during the task—we refer to these electrodes as “single oscillators” and “dual oscillators,” respectively (for example traces see Fig. S1).

Inspecting our data, we observed in many individuals that the frequency of the oscillations at a given hippocampal electrode correlated with its anterior–posterior location, resembling the frequency gradients present in other brain areas (Giocomo & Hasselmo, 2009; Voytek et al., 2010; Zhang et al., 2018). Electrodes at posterior sites often showed oscillations at ~8 Hz. More anterior sites appeared to have oscillations at lower frequencies and more often showed two distinct oscillations (Fig. 3C–D).

**Figure 3:**
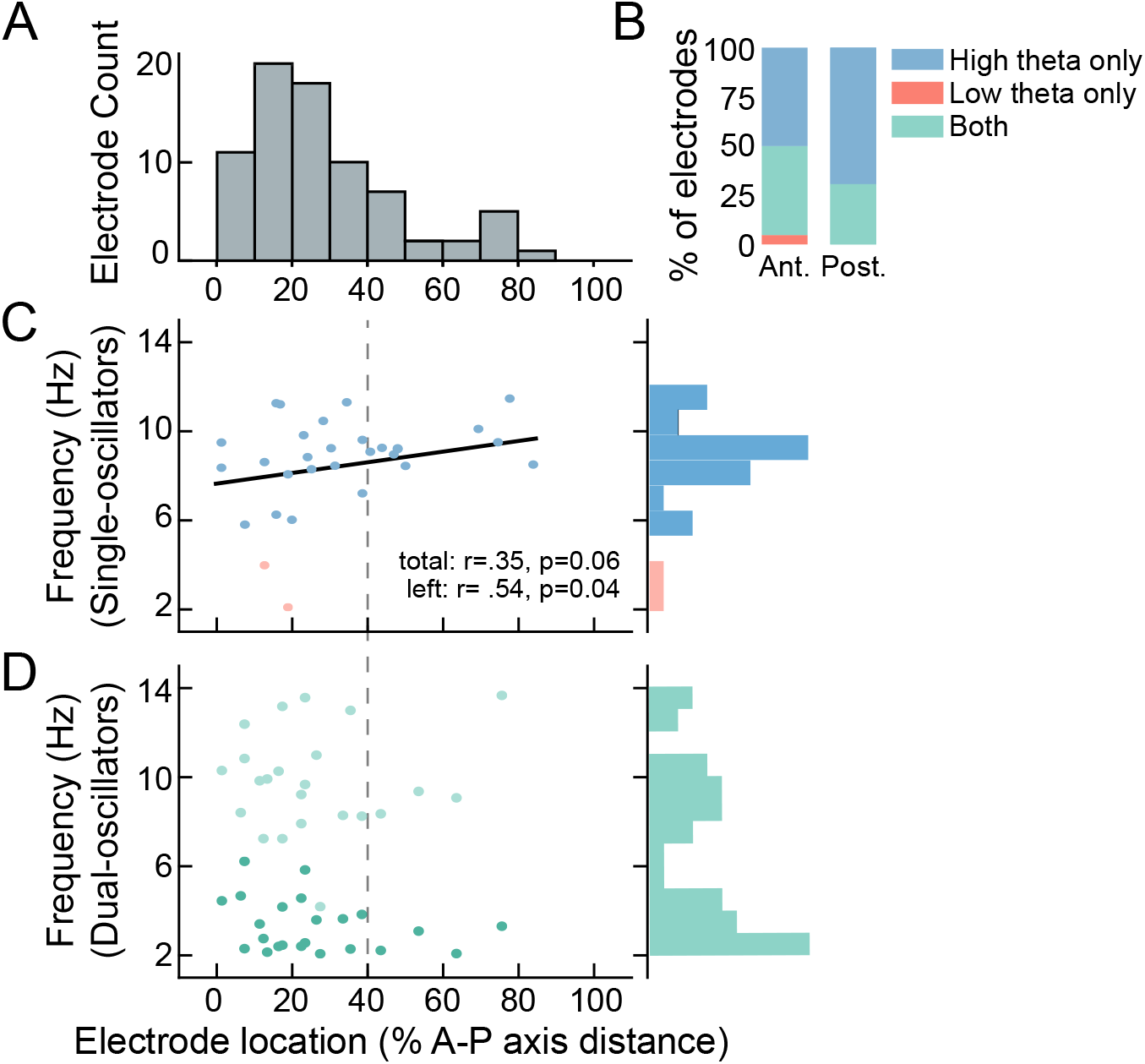
Oscillation properties across frequency and space. **A.** Distribution of electrode locations along the hippocampus anterior–posterior axis. **B.** Proportions of dual oscillators and single oscillators for anterior and posterior hippocampus. **C.** Frequencies and hippocampal localizations of single oscillators across subjects. Black line indicates the fit between frequency and anterior–posterior position for all high-theta electrodes in both hippocampi. Gray dotted line indicates the split between anterior and posterior hippocampus. **D.** Frequency and localization of dual oscillators. Dark green represents the slower oscillation.

We verified these observations quantitatively by analyzing oscillation mean frequencies across our complete dataset. Although individual subjects generally were implanted with only a small number of hippocampal contacts, in aggregate our dataset sampled 80% of the anterior–posterior length of the hippocampus (Fig. 3A). Every hippocampal electrode showed at least one narrowband oscillation within 2–14 Hz (Fig. 3B). 57% (30 of 53) of electrodes were single oscillators, which usually (93%) showed an oscillation in the high-theta (4–14 Hz) band (Fig. 3C). The remaining 43% (23 of 53) of electrodes were dual oscillators (Fig. 3D). In the posterior hippocampus, 69% of electrodes were single oscillators; whereas in the anterior hippocampus, 50% of electrodes were single oscillators and 50% were dual oscillators (Fig. 3B).

These patterns suggested to us that there could be a systematic relationship between the A-P position of a hippocampal recording electrode and the characteristics of the oscillations it exhibited. Indeed, we found that A–P position alone was sufficient to significantly predict whether an electrode was a single or dual oscillator, with single oscillators more prevalent in posterior locations (logistic regression, *p* = 0.02; Fig. 3B). Among the high-theta single oscillators, there was a trend for a gradient between oscillation frequency and anterior–posterior position, such that the specific high-theta frequency of an oscillation was greater for electrodes at more posterior locations (*r* = 0.35, *p* = 0.06; Fig. 3C). This frequency gradient was more clear in the left hippocampus than the right (left: *r* = 0.54, *p* = 0.04; right: *r* = 0.14, *p* = 0.19). Dual oscillators did not show a significant correlation between frequency and location for either the low-or high-theta bands (|*r*| < 0.2, *p*’s > 0.25; Fig. 3D). Together, these results indicate that human hippocampal theta is not a single unitary phenomenon, and that it instead shows a gradient in terms of its features, with more posterior areas behaving more like the theta oscillation seen in rodents by exhibiting a single oscillation at a higher frequency. In contrast, theta oscillations recorded from more anterior locations in the hippocampus were more likely to manifest either as two oscillations or as a single slower oscillation.

Building off our earlier work showing the functional lateralization of human theta oscillations (Miller et al., 2018), here we went further to examine the spectral and anatomical properties of theta oscillations related to movement. We found that both left and right hemispheres displayed low-and high-theta oscillations (29 left electrodes, mean = 6.4 ± 0.5 Hz [mean ± SEM]; 24 right electrodes, mean = 7.1 ± 0.5 Hz). Among the high-theta single oscillators, mean frequencies were significantly faster on the right hemisphere than the left (17 left electrodes; mean = 6.23 Hz, 13 right electrodes; mean = 8.12 Hz; *t*_34_ = 2.4, *p* = 0.02, unpaired *t* test). The high-theta oscillations exhibited by dual oscillators did not differ in frequency between the two hemispheres (*t*_27_ = 0.45, *p* = 0.65, unpaired t-test).

We wished to confirm that our results were not biased by unbalanced electrode positioning across the hippocampus, either between hemispheres or across the dorsal–ventral axis. To analyze this, we compared the distributions of electrode locations across left vs. right hemispheres and across hippocampal subregions (see *Supplemental Results*). We did not find a significant difference in the distribution of electrode positioning between the left vs. right hemispheres (two-sample rank-sum test, *p* = 0.2), and did not find significant frequency variations across subregions (one-way ANOVA; single oscillators: *F*_40_ = 1.8, *p* = 0.18; dual oscillators, *F*_47_ = 0.24, *p* = 0.79). Taken together, these results indicate that the variation in the properties of hippocampal oscillations along the A–P axis is not an artifact of a difference in electrode positioning between the left and right hemispheres or across subregions.

We wished to further probe the spectral characteristics of dual oscillators to understand the relationship between their lower-and higher-theta oscillations. We first considered the possibility that there was a relationship between the particular frequencies of the oscillations that appeared at individual dual oscillator electrodes. This could be the case, for example, if one electrode with two apparent oscillations was actually demonstrating an oscillation and its faster harmonic. However, there was no correlation between the frequencies of the high and low oscillations at individual dual oscillators (p = 0.85, permutation procedure), indicating that the faster oscillations at these sites were not harmonics of the slower ones. Additionally, we compared the timing of the occurrence of these oscillations and found a tendency for dual oscillators to show oscillations at both bands simultaneously (Wilcoxon signed-rank test, *p* = 0.03; see Fig. S2A). Together, these results indicate that signals at dual oscillators reflect distinct hippocampal oscillations with a moderate tendency to co-occur in time. Finally, some subjects within our dataset possessed multiple distantly spaced electrodes along the hippocampal A–P axis. This enabled us to test whether theta oscillations recorded across the A–P axis were temporally related (i.e. through volume conduction). We conducted an analysis of theta phase locking separately for simultaneously recorded electrode pairs that exhibited low-and high-theta oscillations, and found that volume conduction did not explain our findings (see *Supplemental Results*; Fig. S3). Instead, we detected characteristic phase lags indicating that the theta oscillations we were measuring were traveling waves that tended to propagate from higher-theta oscillation sites towards lower-theta ones (Ermentrout & Kleinfeld, 2001; Zhang & Jacobs, 2015; Zhang et al., 2018).

### Analysis of theta-bout duration

Earlier studies showed that theta oscillations in both humans and monkeys appeared in transient bouts (Ekstrom et al., 2005; Watrous, Lee, et al., 2013; Jutras et al., 2013), which were shorter in duration compared to the rodent theta oscillations that often persisted for many seconds (Buzsáki, 2005). To compare our results with signals in rodents, we measured the mean duration of continuous oscillatory cycles of theta signals from individual electrodes in the low-and high-theta bands and for single and dual oscillator electrodes (Fig. 4). Individual electrodes showed a range of mean theta-bout durations. The mean bout duration was longer for high-than for low-theta oscillations (*t*_94_ = 10, *P* < 10^-15^, unpaired t-test). Within the high-theta band, we observed longer theta bouts at single oscillator than dual oscillator electrodes (2.93 vs 2.41 cycles, respectively; Γ68 = 3.3, *p* = 0.002, unpaired t-test). Repeating this analysis in an across-subject fashion yielded similar results (see *Supplemental Results*), confirming that the effects were not driven by individual subjects displaying the effects across multiple electrodes. Overall, although the single oscillator bout durations are still shorter than those seen in rodents (Watrous, Lee, et al., 2013; Jacobs, 2014), their longer high-theta bout durations make them more similar to the theta signals seen in rodents than the shorter bout durations seen in dual oscillators (Buzsáki, 2005).

**Figure 4:**
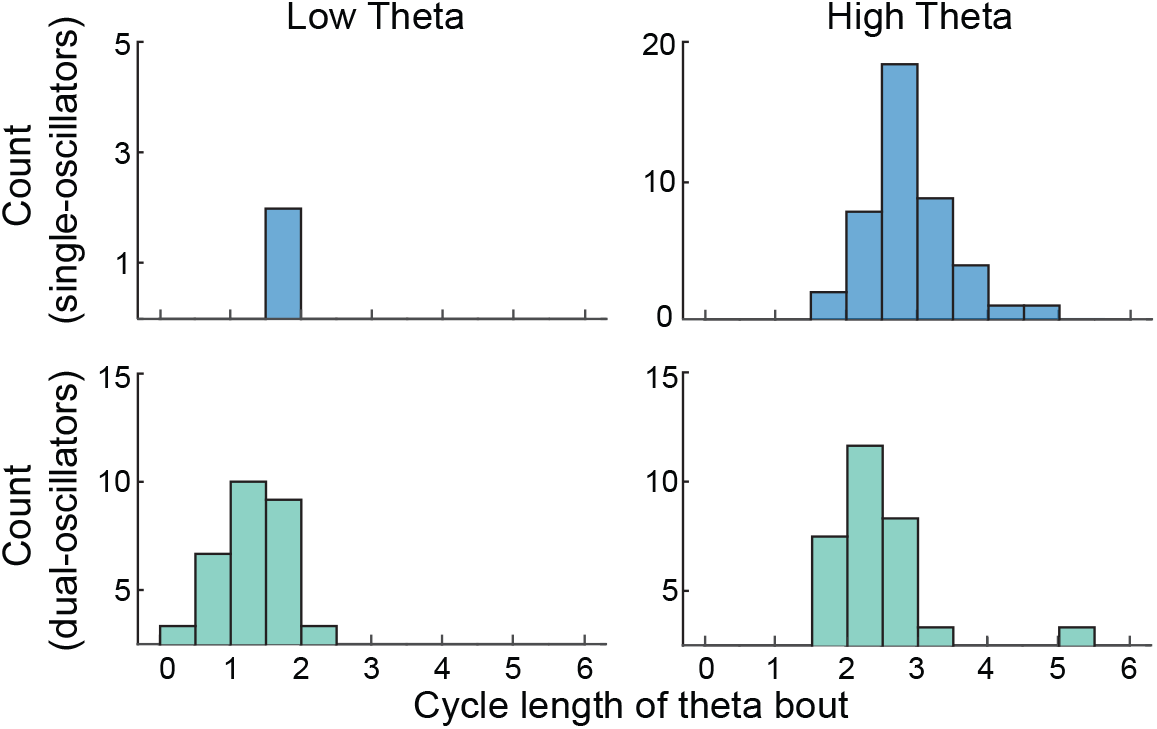
Analysis of the duration of individual theta oscillation bouts. Histograms showing the distributions of mean cycle lengths of the bouts of theta oscillations from individual electrodes. Individual plots show these distributions separately for low-and high-theta oscillations from from single-and dual-oscillator electrodes.

### The frequency of high-theta oscillations correlates with movement speed

In rodents, the instantaneous frequency of the hippocampal theta oscillation correlates with the speed of running (McFarland et al., 1975; Rivas et al., 1996; Geisler et al., 2007; Jeewajee et al., 2008; Hinman et al., 2011; Bender et al., 2015). Further, in both humans and rodents theta power correlates with speed (McFarland et al., 1975; Wyble et al., 2004; Watrous et al., 2011; Hinman et al., 2011). These results have been interpreted to indicate that theta oscillations are important electrophysiological signals that have a general role related to multiple aspects of spatial cognition (Burgess et al., 2007; Jeewajee et al., 2008; Korotkova et al., 2018).

Building off this work, we tested for correlations between movement speed and theta frequency to specifically distinguish the functional role for human hippocampal oscillations in spatial processing. To do this, at each electrode we measured the precise frequency of the oscillations in each of the three movement epochs per trial, in which subjects were moved at a particular fixed speed along the virtual track (see *Methods*). Then, for each electrode, we computed the correlation across epochs between the movement speed and the oscillation frequency.

Many electrodes with high-theta oscillations showed positive correlations between frequency and movement speed. Figure 5A–B illustrates this pattern of results for five example electrodes. We found that the mean correlation between movement speed and oscillation frequency was reliably positive for high-theta oscillations (Fig. 5C, right), both when this signal was observed on single as well as dual oscillator electrodes (both *p*’s < 10^-3^). The mean speed–frequency correlation was significantly larger for single than dual oscillators (*t*_49_ = 4.73, *p* < 10^-4^). This effect was also statistically significant on the single-electrode level. Of the 30 high-theta single oscillators, 24 (86%) showed a significant (*p* < 0.05) speed–frequency correlation, which was more than expected by chance (*p* < 10^-5^, binomial test). Similarly, of 23 dual oscillators, 6 (26%) showed a significant high-theta speed–frequency correlation (p = 0.001, binomial test). We confirmed that these patterns were robust by repeating this analysis after subsampling each trial to contribute only a single randomly selected movement epoch (see *Methods*). The results persisted, with 78% of single oscillators and 44% of dual oscillators showing significant positive correlations (both *p′s* < 0.001).

**Figure 5:**
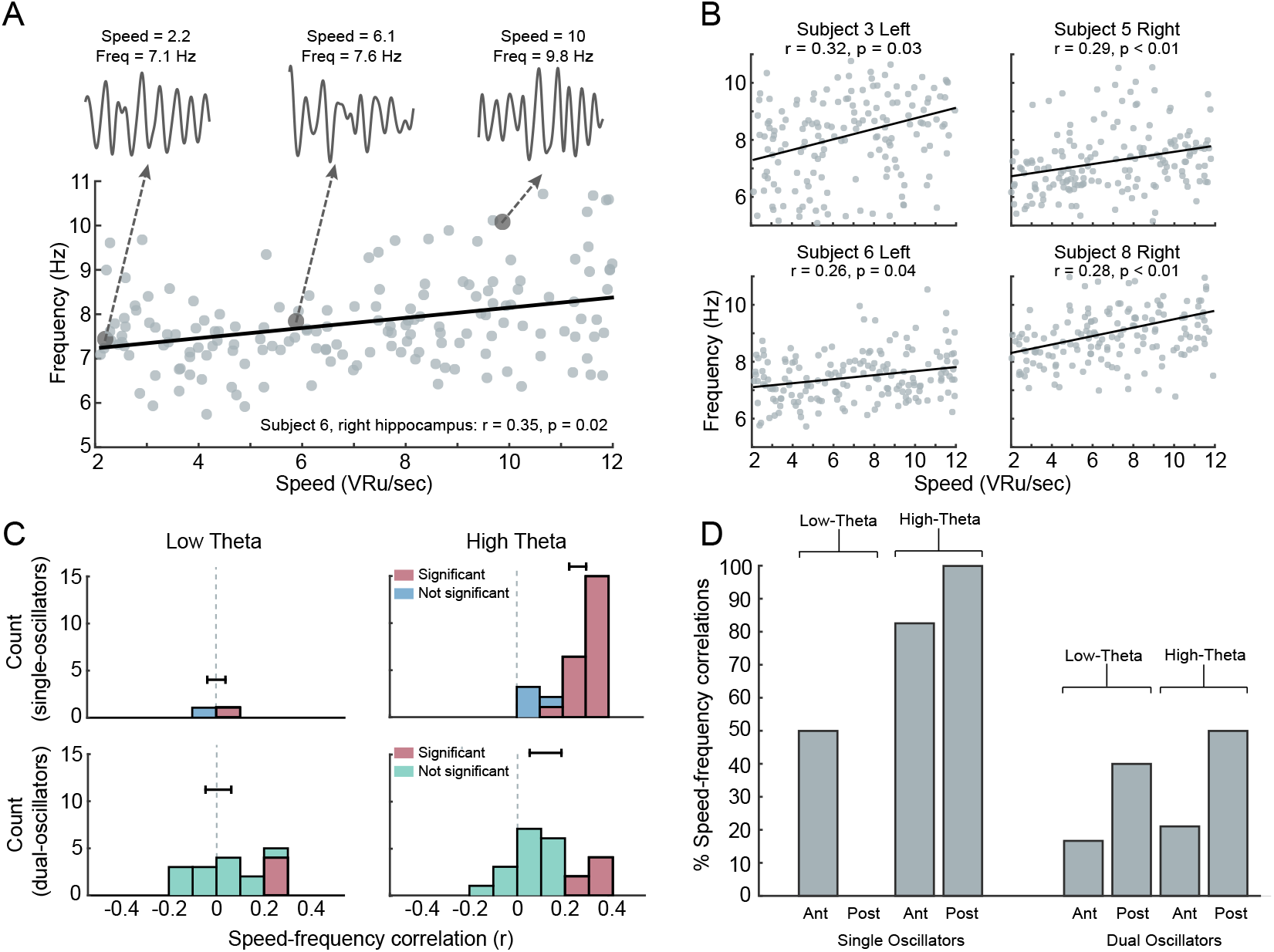
Analyses of the relation between theta frequency and movement speed. **A.** An example electrode with a positive high theta frequency–speed correlation. 2-s trace of filtered hippocampal oscillations during slow, medium, and fast speeds. **B.** Example electrodes from both left and right hippocampus that display significantly positive high theta speed–frequency correlations. **C.** Histogram of correlation coefficients for single and dual oscillators, separately aggregated for low-and high-theta bands. Significant correlations indicated in red. Error bars are SEM. **D.** Percentage of electrodes in each hippocampal region with a significant positive correlation between movement speed and frequency for both low-and high-theta bands.

In contrast to the high-theta band that showed robust positive speed–frequency correlations, in the low-theta band we observed a different pattern, where speed–frequency correlations were not significantly greater than zero (*p′s* > 0.05; Fig. 5C, left). Similarly, at the single-electrode level, of the 25 electrodes with low-theta oscillations (including both single and dual oscillators) only 7 (28%) showed significant speed–frequency correlations, which is significantly less than the proportion of electrodes with high-theta oscillations that showed this effect (z-test, *z* = 2.5, *p* = 0.006). To ensure that these findings were not influenced by individual subjects who displayed the effects strongly at multiple electrodes, we repeated the above analyses in an across-subject fashion (see *Supplemental Results*), and again found that the mean speed–frequency correlation was significantly greater for subjects’ high-theta oscillations compared to their low-theta oscillations (*z* = 2.22, *p* = 0.02).

Finally, given the multiple factors that correlated with theta’s properties—movement speed, electrode location, and oscillation frequency—we performed a multivariate analysis to disentangle the interrelations between these factors. We used a two-way ANOVA to establish how the presence of speed–frequency correlations, averaged within subject, varied across region (anterior/posterior), band (high-/low-theta), and their interaction (Fig 5D). For single oscillators, we found a significant interaction effect (*p* = 0.03) but no significant main effects (A–P region: *p* = 0.09, frequency: *p* = 0.37), thus confirming our interpretation that the prevalence of speed–frequency effects is significantly greater for posteriorly located high-theta single-oscillator electrodes (Fig. 5D). We performed a similar analysis for dual oscillators, and found no significant main or interaction effects (*p′s* > 0.05). This result supports the idea that high-theta oscillations at single oscillators in the posterior hippocampus are specifically involved in spatial processing (Kumaran et al., 2009; Lin et al., 2018).

## Discussion

By recording along the length of the hippocampus from subjects performing a virtual-reality spatial task, we identified multiple patterns of hippocampal theta-band oscillations with separate functional and anatomic properties. These findings suggest that human hippocampal theta oscillations are more than just a slower and noisier analog of the single theta oscillation seen in rodents, but that they instead exist in multiple forms. Specifically, we identified high (~8 Hz) theta oscillations in the posterior hippocampus that varied their frequency with the speed of movement during virtual navigation, similar to the theta oscillations seen in rodents (McFarland et al., 1975; Rivas et al., 1996; Stawińhska & Kasicki, 1998; Jeewajee et al., 2008; Royer et al., 2010; Hinman et al., 2011; Korotkova et al., 2018). We also found that humans have slower hippocampal theta oscillations with distinct functional and anatomical properties. In conjunction with earlier work linking human low-theta to memory (Raghavachari et al., 2001; Lega et al., 2012; Staudigl & Hanslmayr, 2013; Miller et al., 2018), our results suggest that high-and low-theta oscillations represent distinct functional network states. In this way, our work supports the broader view that the brain can exhibit distinct oscillatory states related to different behaviors (Watrous, Tandon, et al., 2013). Further, because the prevalence of spatially relevant single-oscillator electrodes differed along the anterior–posterior length of the hippocampus, our findings provide electrophysiological evidence for a functional gradient across the hippocampus. This is a substantial difference compared to rodents, which usually are described as showing a single theta rhythm across the hippocampus (Lubenov & Siapas, 2009; Long et al., 2015; but see Schmidt et al., 2013).

Previous work on human hippocampal oscillations generally emphasized that rhythms at ~1–5-Hz were more common (for review, see Jacobs, 2014). Our study has several distinctive methodological features to explain why we instead observed a greater prevalence of hippocampal oscillations at faster frequencies. Although not all studies report the intra-hippocampal locations of recording electrodes, it seems that they usually most extensively sampled anterior areas of the hippocampus (e.g. Watrous et al., 2011; Watrous, Lee, et al., 2013). By contrast, our study measured each electrode’s anterior–posterior location and included greater electrode coverage in middle and posterior sections of the hippocampus, which were the regions that more specifically showed single high-theta oscillations. This increased posterior hippocampal coverage is the result of the use of stereotactic electroencephalographic (sEEG) recording electrodes, which have recently grown in popularity for clinical epilepsy mapping (e.g., Lin et al., 2018).

An additional differentiating factor was the design of our behavioral task. Rather than allowing the subject to control their own movement with a fixed top speed as in earlier studies (e.g., Ekstrom et al., 2003; Jacobs et al., 2007, 2013; Miller et al., 2018), here the task automatically changed the subject’s speed randomly, at random times. Given this unpredictable movement, to perform the task well, subjects could not predict their location based on timing and had to continually attend to their view of the spatial environment. We hypothesize that this increased spatial attention increased the prevalence of neural oscillations related to spatial processing. Accordingly, the relatively large prevalence of high-theta oscillations that we observed is consistent with the idea that this signal is particularly important for spatial processing and thus functionally analogous to the “Type 1” theta observed in rodents (Winson, 1978; Bland, 1986; Olvera-Cortés et al., 2004; McNaughton et al., 2006; Burgess, 2008; Korotkova et al., 2018).

Having demonstrated that human slow and fast theta oscillations differ functionally and anatomically, our results raise the question of the functional role of human low theta. One potential explanation is that this signal, which we more often found in the anterior hippocampus, is related to the “Type 2” theta oscillations that had been characterized previously in rodents. Type 2 theta oscillations appear most strongly at ~4 Hz when rodents are stationary and are traditionally linked to anxiety (Kramis et al., 1975; Bland, 1986; Maren et al., 1994; Seidenbecher et al., 2003; Pape et al., 2005; but see Schultheiss et al., 2019). In contrast, current data from humans link oscillations in this low-theta band to memory processing (Lega et al., 2012; Lin et al., 2018; Miller et al., 2018). Therefore, one possibility is that the low-theta oscillations we observed are indeed related to the rodent Type 2 theta, but this signal in humans has a broader functional role beyond anxiety, perhaps including episodic memory and other types of cognitive processes that involve the anterior hippocampus (Bannerman et al., 2004; Mitchell et al., 2008). Our interpretation is consistent with the recent finding that slower, Type 2 theta oscillations in rodents can be generated by a distinct network of cells in the ventral hippocampus (Mikulovic et al., 2018), which is homologous to the anterior hippocampus in humans (Colombo et al., 1988; Amaral & Witter, 1989; Moser & Moser, 1998; Strange et al., 2014).

A notable feature of our findings is identifying many “dual oscillator” electrodes, which seem to reflect hippocampal networks that are capable of exhibiting both low-and high-theta signals. The existence of these dual oscillators may be important theoretically because the hippocampus in both rodents and humans is known to exhibit theta traveling waves that propagate in a posterior-to-anterior (in humans) or dorsal-to-ventral (rodent) direction (Lubenov & Siapas, 2009; Zhang & Jacobs, 2015). One potential mechanism for hippocampal traveling waves is a network of weakly coupled oscillators (Ermentrout & Kleinfeld, 2001; Zhang et al., 2018). The multiple oscillations shown by dual oscillators may reflect the underlying independent oscillators that can lead to the generation of traveling waves when the phase coupling between them is increased. In particular, by showing that oscillatory phase progresses from locations with fast oscillators to those with slower ones (Fig. S3), our results are consistent with the predictions of this model (Ermentrout & Kopell, 1984; Ermentrout & Kleinfeld, 2001; Zhang & Jacobs, 2015; Zhang et al., 2018).

A key result from our work is showing that high-theta oscillations appear in the human hippocampus during movement in virtual reality. Two recent studies measured human hippocampal oscillations from people walking in the physical world and reported high-theta oscillations (Bohbot et al., 2017; Aghajan et al., 2016; but see Meisenhelter et al., 2018). These results were interpreted to suggest that virtual navigation relies on a fundamentally different, lower-frequency oscillatory network state compared to real-world navigation (Yassa, 2018); however, it should be noted that at least one of the studies that previously showed high-theta oscillations in real-world navigation showed examples of these patterns at relatively posterior locations (Bohbot et al., 2017). By demonstrating that humans can show high theta during virtual-reality, our results suggest a different view. We propose that theta oscillations at various frequencies can be prevalent in both virtual and real spatial environments, with the dominant oscillatory frequency being closer to the high-theta range during moments of high spatial demands. It is possible that the earlier VR studies showed relatively less high-theta because their associated tasks required less spatial attention. Overall, both our current findings and this earlier literature lend support to the notion that the human anterior and posterior hippocampi, respectively, are implicated in low-and high-theta oscillations with different behavioral properties (Fanselow & Dong, 2010; Strange et al., 2014).

One reason why theta oscillations are thought to be important functionally is that they coordinate brain-wide networks to synchronize cortical–hippocampal interactions in learning and memory (Siapas et al., 2005; Hyman et al., 2005; Sirota et al., 2008). Therefore, given that we showed that the human hippocampus exhibits two separate theta oscillations in a single task, an important area of future work will be to understand the potential relation of each of these signals to brain-wide neocortical dynamics (von Stein et al., 1999; Libby et al., 2012). In particular, it is notable that the properties of the anterior and posterior hippocampal oscillations resemble the theta and alpha rhythms that are prominent in the frontal and occipital lobes (Voytek et al., 2010; Zhang et al., 2018; Mahjoory et al., 2019), especially including the “midfrontal” theta often found in scalp recordings (Mitchell et al., 2008). Additionally, recent work has shown that local theta oscillations are utilized by the occipital lobe to transiently influence long range fronto-visual dynamics (Han et al., 2019). Given the predominant involvement of the frontal and occipital lobes in high-level and sensory processing, respectively, this suggests that low-and high-theta oscillations could correlate with different types of hippocampal–neocortical interactions related to distinct functional processes (Libby et al., 2012; Watrous, Tandon, et al., 2013). This multiplicity of human theta patterns could allow the human hippocampus to coordinate a diverse set of brain-wide neural assemblies to support various types of behaviors, including spatial navigation, memory, and other cognitive processes (Kahana et al., 2001; Buzsaki & Moser, 2013; Eichenbaum & Cohen, 2014).

## Methods

### Subjects

Fourteen subjects (8 males and 6 females, age range 23-49) at four hospitals (Thomas Jefferson University Hospital, Columbia University Medical Center, University of Texas Southwestern Medical Center, and Emory University Hospital) undergoing treatment for medication-resistant epilepsy participated in our study. Neurosurgeons implanted these subjects with clinical depth electrodes for functional mapping and the localization of seizure foci. Implantation sites were determined solely by clinical teams, though electrodes were often placed in medial temporal lobe regions that are of interest experimentally. Research protocols were approved by the institutional review boards at each participating hospital, and informed consent was obtained from all subjects. Previous work utilizing scalp EEG recordings (Kober & Neuper, 2011; Ramos-Loyo & Sanchez-Loyo, 2011) has reported that theta oscillatory activity varied with age and sex. However, here we did not find a significant relation between theta frequency and age (*r* = −0.09, *p* = 0.90) or sex (*t*_11_ = 0.97, *p* = 0.35, unpaired t-test). This difference suggests that the hippocampal oscillations that are the focus of our study differ from the neocortical signals measured with scalp EEG.

### Task

The subjects in our study performed a new spatial memory task, which we specifically designed to encourage subjects to pay attention to their location in the virtual environment by varying their movement speed randomly. This distinctive design prevented subjects from utilizing a timing-based strategy to perform the task, such as by by remembering each object’s latency since the beginning of movement. We hypothesized that this task design had the potential to elicit more reliable hippocampal activity related to spatial processing than previous studies of human navigation (Qasim et al., 2019). Because the subjects in our study were undergoing continuous monitoring for epileptiform activity, we were limited to studying virtual navigation, as subjects remained in their hospital bed throughout testing.

In the task, subjects were moved along the length of a virtual reality (VR) track, which we defined as having a length of 70 virtual reality units. The ground was textured to mimic asphalt, and the track was surrounded by stone walls (see Fig. 1). On each trial, subjects were placed at the beginning of the track, and they began each trial by pressing a button on a game controller. Next, a four-second-long countdown timer appeared. After the countdown, subjects were moved forward along the track. Within each third of the track, subjects were moved at a constant speed, which was chosen randomly from a uniform distribution between 2 and 12 VR-units/second. Locations where speed changes began are indicated by the light gray shading in the schematic shown in Figure 1C. When speed changes occurred, acceleration occurred gradually over the course of one second to avoid jarring transitions.

During movement, the subjects’ goal was to mark the location of a hidden object. The first two times that the subject traveled down the track, the object’s location was visible (Fig. 1A). On subsequent trials, the object was invisible, and subjects were instructed to press the button on the controller when they believed they were at the correct location (Fig. 1B). The closer the subject pressed the button to the correct location, the more points they received (as indicated in the top right of the display), thus encouraging careful attention to current location in the environment. Subjects were also required to press the button when they approached the end of the track where the ground was colored red to ensure that they were attentive during the trial. Possible object locations are indicated by the dark gray shading in Figure 1C.

Each trial consisted of the subject traveling a single time down the track, either encoding or retrieving object location. Within each trial, the task would automatically change the subject’s speed at each of two possible speed-change regions (Fig. 1C), such that the subject’s path down the track consisted of three constant speed regions. The focus of this study was to analyze human hippocampal correlates of movement, rather than memory or task performance. Thus, we analyzed all time points while subjects were in motion, regardless of performance, including both encoding and retrieval trials. We classified “correct” trials as trials where subjects had a response–object error distance of less than 11.5 VR units. This value represents the average error that subjects would display if they responded at the midpoint of the track each time (Qasim et al., 2019).

### Electrophysiology

We recorded subjects’ intracranial electroencephalographic (iEEG) data from implanted depth electrodes via the clinical or research recording systems present at the participating hospitals (Nihon Kohden; XLTEK; Neuralynx; Blackrock). Data were recorded at a sampling rate of either 1000 or 2000 Hz. iEEG signals were initially referenced to common intracranial or scalp contacts, and were subsequently re-referenced using an anatomically weighted referencing scheme prior to analysis. Data were notch filtered at 60 Hz using a zero-phase-distortion Butterworth filter to remove line noise prior to subsequent analyses. iEEG recordings were aligned to the behavioral task laptop via synchronization pulses sent to the recording system.

### Electrode Localization

Our data analyses were designed to test how the functional and electrophysiological properties of human theta oscillations varied along the hippocampal anterior–posterior (A–P) axis. To study electrode’s anatomical features, we localized depth electrodes for each subject using an established semi-automated image processing pipeline (Jacobs et al., 2016). We delineated the hippocampus, we applied the Automatic Segmentation of Hippocampal Subfields multi atlas segmentation method to pre-implantation high-resolution hippocampal coronal T2-weighted and whole-brain 3D T1-weighted scans. Electrode contact coordinates derived from post-implantation CT scans were then co-registered to the segmented MRI scans using Advanced Normalization Tools (Avants et al., 2008), and anatomic locations were automatically generated. A neuroradiologist reviewed and confirmed contact locations based on the co-registered source images. Electrodes were assigned normalized locations along the hippocampal axis by determining the coronal slice containing the center of the contact and measuring relative to the first and last MRI slice containing the hippocampus. For specific subjects, a neuroradiologist generated transparent 3D surface renderings of the subjects’ hippocampal segmentation and corresponding co-registered electrode contacts. Here we only analyzed electrodes located within the hippocampal formation (CA1, CA2, subiculum, and dentate gyrus). For the majority of our analyses, we analyzed electrode location as a continuous variable along the hippocampal A–P axis; however, when it was more convenient to refer to “anterior” and “posterior” labeling, we utilized 40% as the division point, based on the midpoint of our coverage, to allow adequate statistical power for data analyses. If two or more neighboring electrodes in one subject were located in nearby slices (less than 10% of the hippocampal A-P axis distance away from each other), and exhibited a similar oscillation frequency (within 2 Hz) during movement, all but one of these electrodes were dropped for all analyses.

### Spectral Analysis

Due to the variability of human neuronal oscillations (Lega et al., 2012; Zhang & Jacobs, 2015), our analyses examined the spectral features of the oscillations at each electrode at a high resolution to identify frequency bands that are customized for each subject and electrode (Manning et al., 2009; Zhang et al., 2018). This approach differs from the one used in our earlier work, which utilized fixed frequency bands across subjects. To achieve this high frequency resolution, we followed the MODAL algorithm (Watrous et al., 2018). The first step of this algorithm is to exclude epochs of the data that could potentially result from epileptic activity (Gelinas et al., 2016). Then, the algorithm defines relevant frequency bands as those frequencies where the measured oscillatory power exceeds one standard deviation above the background 1/*f* spectrum. This criterion ensures that our results were not driven by spurious background noise information. MODAL then computes the instantaneous frequency and phase for each frequency band, but only when the local power spectrum (computed in 10 second, non-overlapping windows) indicated a local power peak that band.

When examining frequency band characteristics across the data, we noticed that every electrode exhibited either one or two distinct oscillations at frequency bands between 2 and 14 Hz. We called electrodes that only exhibited a single oscillation throughout the task “single oscillators” while we called those that exhibited two oscillations “dual oscillators.” For an electrode to be designated as a dual oscillator, the edges of the two frequency bands detected by MODAL had to differ by least 0.5 Hz. For analyses where we specifically report low-theta and high-theta results across electrodes, we classified “low-theta” oscillations as those < 4 Hz, and “high-theta” oscillations as those >= 4 Hz. We defined an ‘‘oscillatory bout” as sequences of consecutive millisecond time points of any length where at least one oscillation was present.

We performed a series of analyses comparing how these detected oscillations related to features of the subject’s movement. Each trial within the task consisted of three intervals that each had a constant speed of movement (Fig. 1). We computed the particular oscillation frequency for each movement interval by first using MODAL to measure the instantaneous frequency of the iEEG signal at each timepoint throughout the interval. Then, we computed a histogram of the distribution of frequencies (0.1-Hz bins), identified the single most-often occurring frequency (i.e., the mode), and used this value to summarize the oscillatory activity in that interval. For our initial analysis of speed-frequency correlations (Fig 5A-C), we utilized all three speeds from each trial to compute three speed–frequency correlations per trial. To ensure independence, we also repeated the analysis by randomly choosing only a single speed period from each trial to analyze, which produced comparable results.

## Acknowledgements

This work was supported by the National Institutes of Health (R01-MH104606, S10-OD018211), and the National Science Foundation (Graduate Research Fellowship DGE 16-44869). We thank Shachar Maidenbaum for providing thoughtful comments on the manuscript.

## Supplementary Results

### Analysis of Hippocampal Subregions

We considered the possibility that our results could have been affected by other anatomical factors besides A–P location. To test for an effect of the electrode’s dorsal–ventral (D–V) position, we compared the relation between the D-V location (calculated for each electrode in the same manner as A-P location - see *Methods*) of each electrode within the hippocampus and the frequency of the measured oscillations, both for single and dual oscillators. We did not find a significant relation between frequency and D-V location for low-or high-theta oscillations for either single or dual oscillators (all p’s > 0.05). To further probe potential effects of electrode positioning, we performed a new analysis where we manually identified each electrode’s subregion. Then, to aggregate data across subjects, we compared signals between these subregions (CA1, CA2, DG, and subiculum). This procedure allowed us to examine potential effects of D-V location while accounting for inter-individual morphological differences. For both single and dual oscillators, theta frequency did not significantly change across subregion (single oscillators: *F*_40_ = 1.8, *p* = 0.18; dual oscillators, *F*_47_ = 0.24, *p* = 0.79). Together, these results indicate that our primary results of theta frequency shifting with A–P position (Fig. 3) are not confounded by variations related to subregion or D–V location.

### Group-level Analyses

For the analyses of oscillatory bouts (Fig. 4) and frequency–speed correlations (Fig. 5), we wished to ensure that our results were not influenced by subjects who had multiple electrodes at similar A–P locations. Therefore, we performed a group-level analysis where each subject contributed only a single mean value per frequency/region, for each of the low-anterior, low-posterior, high-anterior, and high-posterior categories. Thus, in this analysis each subject contributed at most one datapoint per category. The results of this analysis recapitulated our main findings. The mean number of cycles for low-theta oscillations was significantly shorter than that of high-theta oscillations (1.58 vs. 2.87 cycles, respectively; *t*(38) = 12.0, *p* < 10^-13^). Analogously, the mean number of cycles for high-theta oscillations in single oscillators is longer compared to high-theta oscillations in dual oscillators (3.0 vs. 2.43 cycles, respectively; *t*(25) = 6.0, *p* < 10^-5^). Similarly, across subjects, we found that the mean speed–frequency correlation was significantly greater for a subject’s high-theta oscillations compared to his or her low-theta oscillations (*z* = 2.22, *p* = 0.02).

### Timing analysis between dual oscillator bouts

We characterized the temporal relationship between the two oscillations that appeared on dual oscillator electrodes during movement to determine whether they tended to occur simultaneously or at alternate timepoints (Fig. S2). For this analysis, we identified all oscillations on each band for dual oscillators that were at least 2 cycles in length. Then, for each electrode, we labeled each timepoint with two binary variables, indicating the oscillation’s presence or absence for each band. Finally, for each electrode, we computed the *ϕ* correlation coefficient between the binary variables for each of the two bands—a positive *ϕ* would indicate that an electrode tended to exhibit oscillations at both bands simultaneously. Comparing the distribution of *ϕ* values across electrodes, we found the mean *ϕ* was significantly positive (mean *ϕ* = 0.07, Wilcoxon signed-rank test, *p* = 0.03; Fig. S2A). This result indicates that dual oscillators tended to show oscillations at both of its bands simultaneously.

### Low-and High-Theta Phase Analysis

We wanted to understand the spatial organization of theta hippocampal oscillations at different frequencies along the A–P axis, inspired by our earlier finding of theta traveling waves that propagate from regions with fast to slow relative frequencies (Zhang et al., 2018). To probe this issue, we identified data from subjects who had pairs of simultaneously implanted electrodes spanning at least 30% of their hippocampus. We then analyzed the instantaneous phase at each electrode and computed the mean phase difference between those electrodes in each trial. We then tested whether the theta phases from each electrode in the pair were correlated using circular statistics (Berens, 2009). The results were as follows: 21% of high-theta electrode pairs showed theta oscillations with independent phase patterns (Rayleigh test for phase uniformity, *p* > 0.05). 45% of pairs showed high-theta oscillations that were synchronized (Rayleigh test *p* < 0.05) with a 0° ± 10° phase shift—this indicates that synchronized theta oscillations appeared in both electrodes with the same timing, as predicted by volume conduction. 33% of electrode pairs showed high-theta oscillations with a consistent phase shift between electrodes, which is suggestive of a traveling wave.

We examined the propagation patterns from the electrode pairs with consistent phase shifts to test a prediction of the weakly-coupled-oscillator model (Ermentrout & Kleinfeld, 2001), that traveling waves would propagate towards regions with slower oscillation frequency (Zhang et al., 2018). We tested for this pattern in our data by measuring the sign of the phase shift across the electrode pair and testing whether the sign of this phase shift correlated with the electrodes’ difference in relative frequency. Consistent with the coupled oscillator model, we found that the direction of traveling wave propagation followed the orientation of local frequency gradients. For the pairs where the more posterior electrode showed a faster oscillation frequency, 65% of the time, the pair showed traveling waves propagating in a posterior-to-anterior direction (Fig. S3B–C). This directionality was less consistent for the electrode pairs that had a frequency gradient with the reverse orientation (χ^2^ = 3.4, *p* = 0.06), indicating that having posterior-to-anterior traveling theta waves is correlated with having faster oscillations towards the posterior hippocampus.

---

**Figure S1:**
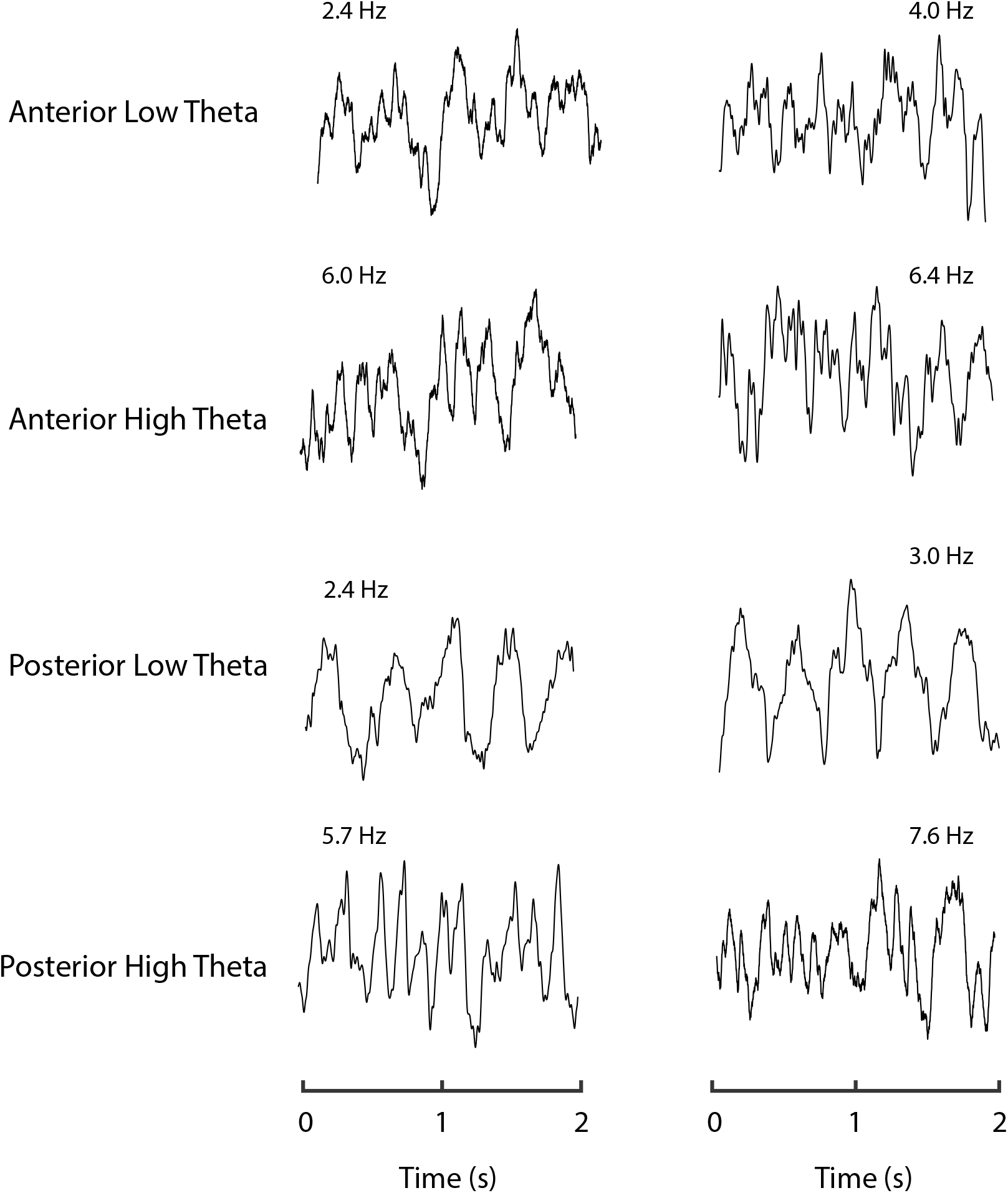
Raw EEG Traces. Raw EEG traces from different subjects within our dataset. Data were lowpass filtered at 55 Hz to filter out line-noise.

**Figure S2:**
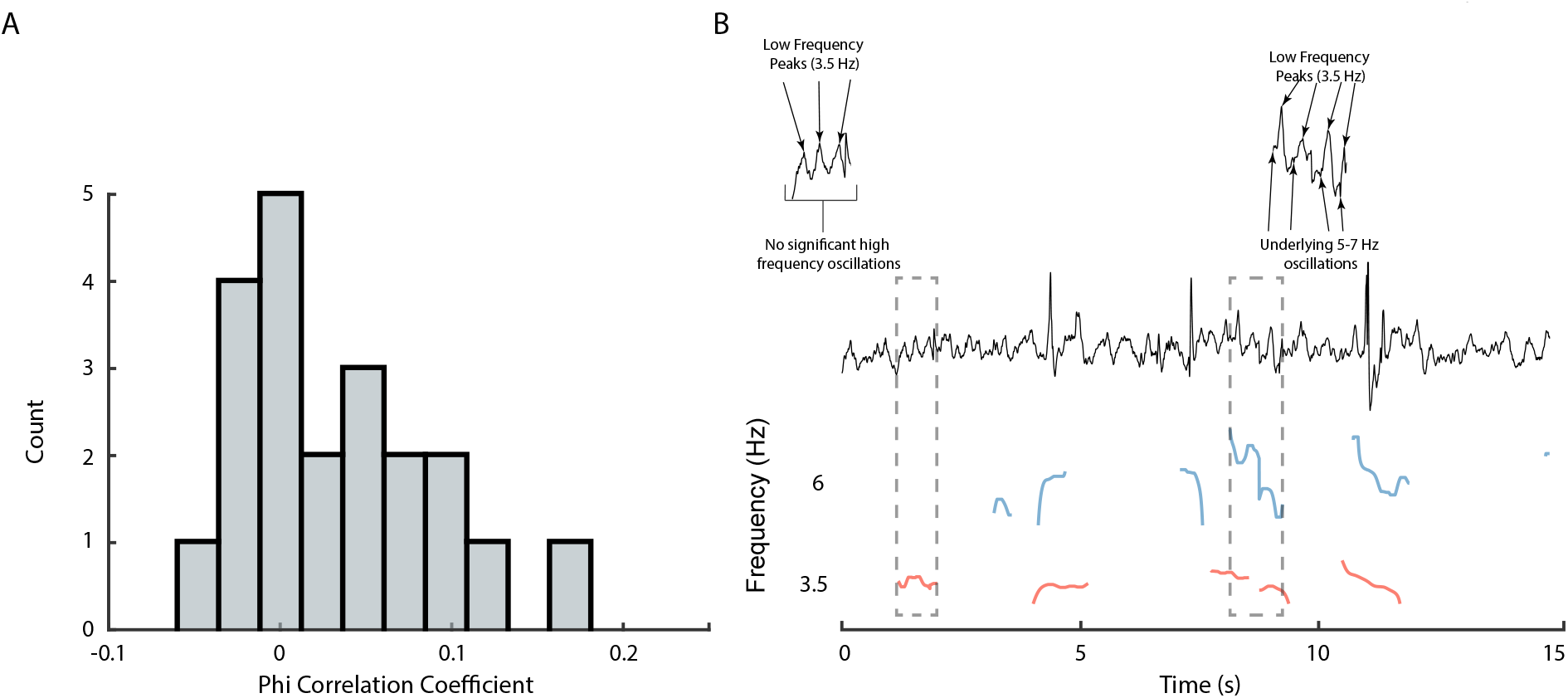
Dual Oscillator Bout Correlation. **A.** Histogram of *ϕ* correlation coefficients from all dual oscillators within our dataset. The distribution is significantly greater than 0 (Wilcoxon signed-rank test, *p* = 0.03). **B.** Raw EEG trace above high-theta and low-theta prevalences for a random 15s from an electrode within our dataset that exhibits characteristic periods where the low-and high-theta bouts appear separately (left) and co-occur (right).

**Figure S3:**
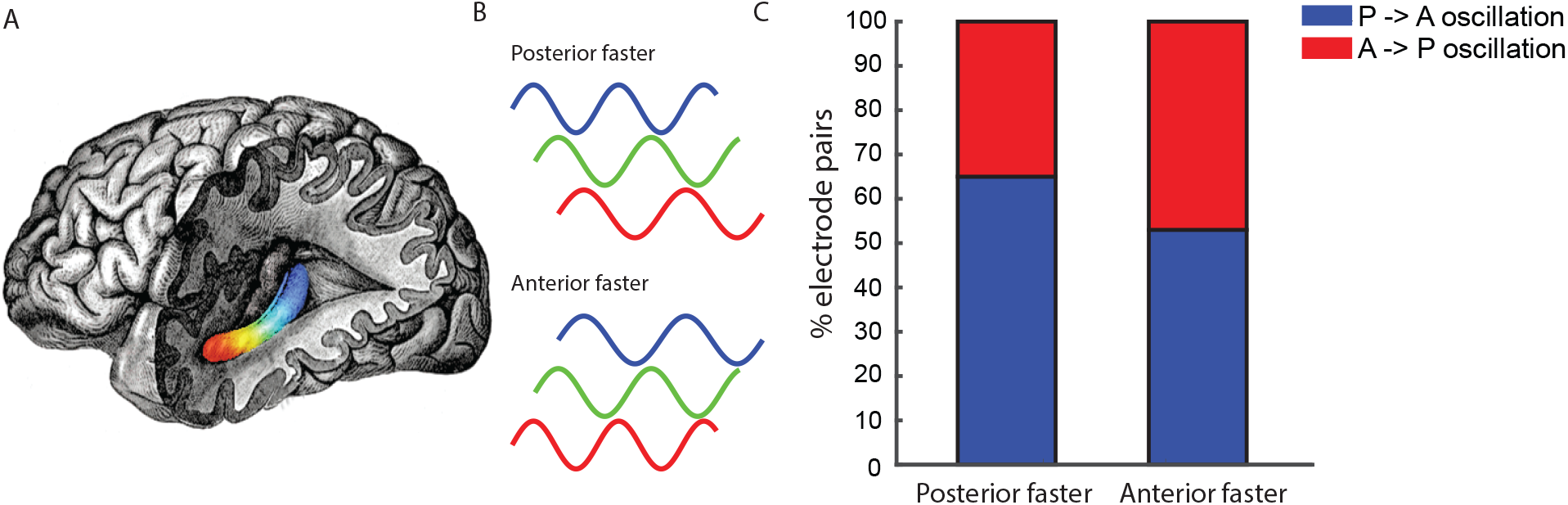
Phase Analysis. **A.** Schematic of the human hippocampus, with colors signifying location along the A-P axis (red is anterior, blue is posterior). **B.** Example iEEG traces from electrodes within a single subject at similar frequencies demonstrating phase shifts (not real data). **C.** For all electrode pairs that exhibit consistent phase shifts, the plot indicates the percent of electrode pairs that exhibit anterior-to-posterior traveling waves (red) and posterior-to-anterior traveling waves (blue), with electrode pairs separated by the site that showed the faster oscillation.

## References

Aghajan, Z. M., Schuette, P., Fields, T., Tran, M., Siddiqui, S., Hasulak, N.… others (2016). Theta oscillations in the human medial temporal lobe during real world ambulatory movement. Current Biology, 27(24), 3743–3751.

Amaral, D. G., & Witter, M. P. (1989). The three-dimensional organization of the hippocampal formation: a review of anatomical data. Neuroscience, 31(3), 571–591.

Arnolds, D. E. A. T., Lopes Da Silva, F. H., Aitink, J. W., Kamp, A., & Boeijinga, P. (1980). The spectral properties of hippocampal EEG related to behaviour in man. Electroencephalography and Clinical Neurophysiology, 50, 324–328.

Avants, B. B., Epstein, C. L., Grossman, M., & Gee, J. C. (2008). Symmetric diffeomorphic image registration with cross-correlation: evaluating automated labeling of elderly and neurodegenerative brain. Medical Image Analysis, 12(1), 26–41.

Bannerman, D., Rawlins, J., McHugh, S., Deacon, R., Yee, B., Bast, T., … Feldon, J. (2004). Regional dissociations within the hippocampus–memory and anxiety. Neuroscience & Biobehavioral Reviews, 28(3), 273–283.

Bender, F., Gorbati, M., Cadavieco, M. C., Denisova, N., Gao, X., Holman, C., … Ponomarenko, A. (2015). Theta oscillations regulate the speed of locomotion via a hippocampus to lateral septum pathway. Nature communications, 6, 8521.

Berens, P. (2009). Circstat: A matlab toolbox for circular statistics. Journal of Statistical Software, 31(10).

Bland, B. H. (1986). The physiology and pharmacology of hippocampal formation theta rhythms. Prog. Neurobiol., 26, 1–54.

Bohbot, V. D., Copara, M. S., Gotman, J., & Ekstrom, A. D. (2017). Low-frequency theta oscillations in the human hippocampus during real-world and virtual navigation. Nature Communications, 8, 14415.

Burgess, N. (2008). Grid cells and theta as oscillatory interference: Theory and predictions. Hippocampus, 18, 1157–1174.

Burgess, N., Barry, C., O’Keefe, J., & London, U. (2007). An oscillatory interference model of grid cell firing. Hippocampus, 17(9), 801–12.

Bush, D., Bisby, J. A., Bird, C. M., Gollwitzer, S., Rodionov, R., Diehl, B., … Burgess, N. (2017). Human hippocampal theta power indicates movement onset and distance travelled. Proceedings of the National Academy of Science USA, 114(46), 12297–12302.

Buzsáki, G. (2005). Theta rhythm of navigation: Link between path integration and landmark navigation, episodic and semantic memory. Hippocampus, 15, 827–840.

Buzsaki, G., & Moser, E. (2013). Memory, navigation and theta rhythm in the hippocampal-entorhinal system. Nature Neuroscience, 16(2), 130–138.

Colombo, M., Fernandez, T., Nakamura, K., & Gross, C. (1988). Functional differentiation along the anterior-posterior axis of the hippocampus in monkeys. Journal of Neurophysiology, 80(2), 1002–1005.

Eichenbaum, H., & Cohen, N. J. (2014). Can we reconcile the declarative memory and spatial navigation views on hippocampal function? Neuron, 83(4), 764–770.

Ekstrom, A. D., Caplan, J., Ho, E., Shattuck, K., Fried, I., & Kahana, M. (2005). Human hippocampal theta activity during virtual navigation. Hippocampus, 15, 881–889.

Ekstrom, A. D., Kahana, M. J., Caplan, J. B., Fields, T. A., Isham, E. A., Newman, E. L., & Fried, I. (2003). Cellular networks underlying human spatial navigation. Nature, 425, 184–187.

Ermentrout, G., & Kleinfeld, D. (2001). Traveling Electrical Waves in Cortex Insights from Phase Dynamics and Speculation on a Computational Role. Neuron, 29(1), 33–44.

Ermentrout, G., & Kopell, N. (1984). Frequency plateaus in a chain of weakly coupled oscillators,. SIAM Journal on Mathematical Analysis, 15(2), 215–237.

Fanselow, M. S., & Dong, H.-W. (2010). Are the dorsal and ventral hippocampus functionally distinct structures? Neuron, 65(1), 7–19.

Geisler, C., Robbe, D., Zugaro, M., Sirota, A., & Buzsaki, G. (2007). Hippocampal place cell assemblies are speed-controlled oscillators. Proceedings of the National Academy of Sciences, USA, 104(19), 8149.

Gelinas, J. N., Khodagholy, D., Thesen, T., Devinsky, O., & Buzsaki, G. (2016). Interictal epileptiform discharges induce hippocampal–cortical coupling in temporal lobe epilepsy. Nature medicine, 22(6), 641.

Giocomo, L. M., & Hasselmo, M. E. (2009). Knock-out of hcn1 subunit flattens dorsal-ventral frequency gradient of medial entorhinal neurons in adult mice. Journal of Neuroscience, 29(23), 7625–7630.

Greicius, M. D., Krasnow, B., Boyett-Anderson, J. M., Eliez, S., Schatzberg, A. F., Reiss, A. L., & Menon, V. (2002). Regional analysis of hippocampal activation during memory encoding and retrieval: fmri study regional analysis of hippocampal activation during memory encoding and retrieval: an fmri study. Hippocampus, 13(1), 164–174.

Han, H.-B., Lee, K. E., & Choi, J. H. (2019). Functional dissociation of theta oscillations in the frontal and visual cortices and their long-range network during sustained attention. eNeuro, 6(6).

Hinman, J. R., Penley, S. C., Long, L. L., Escabi, M. A., & Chrobak, J. J. (2011). Septotemporal variation in dynamics of theta: speed and habituation. Journal of Neurophysiology, 105(6), 2675–2686.

Hyman, J., Zilli, E., Paley, A., & Hasselmo, M. (2005). Medial prefrontal cortex cells show dynamic modulation with the hippocampal theta rhythm dependent on behavior. Hippocampus, 15(6), 739–749.

Jacobs, J. (2014). Hippocampal theta oscillations are slower in humans than in rodents: implications for models of spatial navigation and memory. Philosophical Transactions of the Royal Society B: Biological Sciences, 369(1635), 20130304.

Jacobs, J., Kahana, M. J., Ekstrom, A. D., & Fried, I. (2007). Brain oscillations control timing of single-neuron activity in humans. Journal of Neuroscience, 27(14), 3839–3844.

Jacobs, J., Miller, J., Lee, S. A., Coffey, T., Watrous, A. J., Sperling, M. R., … Rizzuto, D. S. (2016, December). Direct electrical stimulation of the human entorhinal region and hippocampus impairs memory. Neuron, 92(5), 1–8.

Jacobs, J., Weidemann, C. T., Miller, J. F., Solway, A., Burke, J. F., Wei, X., … Kahana, M. J. (2013). Direct recordings of grid-like neuronal activity in human spatial navigation. Nature Neuroscience, 16(9), 1188–1190. doi: 10.1038/nn.3466

Jeewajee, A., Barry, C., O’Keefe, J., & Burgess, N. (2008). Grid cells and theta as oscillatory interference: electrophysiological data from freely moving rats. Hippocampus, 18(12), 1175–1185.

Jutras, M. J., Fries, P., & Buffalo, E. A. (2013). Oscillatory activity in the monkey hippocampus during visual exploration and memory formation. Proceedings of the National Academy of Sciences, USA, 110(32), 13144–13149.

Kahana, M. J., Seelig, D., & Madsen, J. R. (2001). Theta returns. Current Opinion in Neurobiology, 11, 739–744.

Kober, S. E., & Neuper, C. (2011). Sex differences in human eeg theta oscillations during spatial navigation in virtual reality. International Journal of Psychophysiology, 79(3), 347–355.

Korotkova, T., Ponomarenko, A., Monaghan, C. K., Poulter, S. L., Cacucci, F., Wills, T., … Lever, C. (2018). Reconciling the different faces of hippocampal theta: the role of theta oscillations in cognitive, emotional and innate behaviors. Neuroscience & Biobehavioral Reviews, 85, 65–80.

Kramis, R., Vanderwolf, C., & Bland, B. (1975). Two types of hippocampal rhythmical slow activity in both the rabbit and the rat: relations to behavior and effects of atropine, diethyl ether, urethane, and pentobarbital. Exp Neurol, 49(1 Pt 1), 58–85.

Kumaran, D., Summerfield, J. J., Hassabis, D., & Maguire, E. A. (2009). Tracking the emergence of conceptual knowledge during human decision making. Neuron, 63(6), 889–901.

Lega, B., Jacobs, J., & Kahana, M. (2012). Human hippocampal theta oscillations and the formation of episodic memories. Hippocampus, 22(4), 748–761.

Lega, B., Kahana, M. J., Jaggi, J. L., Baltuch, G. H., & Zaghloul, K. A. (2011). Neuronal and oscillatory activity during reward processing in the human ventral striatum. NeuroReport, 22(16), 795–800.

Libby, L. A., Ekstrom, A. D., Ragland, D., & Ranganath, C. (2012). Differential connectivity of perirhinal and parahippocampal cortices within human hippocampal subregions revealed by high-resolution functional imaging. Journal of Neuroscience, 19, 6550–6560.

Lin, J.-J., Umbach, G., Rugg, M. D., & Lega, B. (2018). Gamma oscillations during episodic memory processing provide evidence for functional specialization in the longitudinal axis of the human hippocampus. Hippocampus, 29(2), 68–72.

Long, L. L., Bunce, J. G., & Chrobak, J. J. (2015). Theta variation and spatiotemporal scaling along the septotemporal axis of the hippocampus. Frontiers in systems neuroscience, 9, 37.

Lubenov, E. V., & Siapas, A. G. (2009). Hippocampal theta oscillations are travelling waves. Nature, 459(7246), 534–539.

Maguire, E., Frackowiak, S. J., & Frith, C. D. (1997). Recalling routes around london: activation of the right hippocampus in taxi drivers. Journal of Neuroscience, 17, 7103–7110.

Mahjoory, K., Schoffelen, J.-M., Keitel, A., & Gross, J. (2019). The frequency gradient of human resting state brain oscillations follows cortical hierarchies. bioRxiv.

Manning, J. R., Jacobs, J., Fried, I., & Kahana, M. J. (2009). Broadband shifts in local field potential power spectra are correlated with single-neuron spiking in humans. Journal of Neuroscience, 29(43), 13613–13620. doi: 10.1523/JNEUROSCI.2041-09.2009

Maren, S., DeCola, J. P., Swain, R. A., Fanselow, M. S., & Thompson, R. F. (1994). Parallel augmentation of hippocampal long-term potentiation, theta rhythm, and contextual fear conditioning in water-deprived rats. Behavioral neuroscience, 108(1), 44.

McFarland, W. L., Teitelbaum, H., & Hedges, E. K. (1975). Relationship between hippocampal theta activity and running speed in the rat. Journal of comparative and physiological psychology, 88(1), 324.

McNaughton, N., Ruan, M., & Woodnorth, M.-A. (2006). Restoring theta-like rhythmicity in rats restores initial learning in the morris water maze. Hippocampus, 16(12), 1102–1110.

Meisenhelter, S., Testorf, M. E., Gorenstein, M. A., Hasulak, N. R., Tcheng, T. K., Aronson, J. P., & Jobst, B. C. (2018). Cognitive tasks and human ambulatory electrocorticography using the rns system. Journal of neuroscience methods, 311, 408–417.

Mikulovic, S., Restrepo, C. E., Siwani, S., Bauer, P., Pupe, S., Tort, A. B., … Leao, R. N. (2018). Ventral hippocampal olm cells control type 2 theta oscillations and response to predator odor. Nature Communications, 9(1), 3638.

Miller, J. F., Watrous, A. J., Tsitsiklis, M., Lee, S. A., Sheth, S. A., Schevon, C. A.,… Jacobs, J. (2018). Lateralized hippocampal oscillations underlie distinct aspects of human spatial memory and navigation. Nature communications, 9(1), 2423.

Mitchell, D., McNaughton, N., Flanagan, D., & Kirk, I. (2008). Frontal-midline theta from the perspective of hippocampal theta. Progress in neurobiology, 86(3), 156.

Moser, M., & Moser, E. (1998). Functional differentiation in the hippocampus. Hippocampus, 8(6), 608–619.

Olvera-Cortes, E., Guevara, M., & González-Burgos, I. (2004). Increase of the hippocampal theta activity in the Morris water maze reflects learning rather than motor activity. Brain research bulletin, 62(5), 379–384.

Pape, H.-C., Narayanan, R. T., Smid, J., Stork, O., & Seidenbecher, T. (2005). Theta activity in neurons and networks of the amygdala related to long-term fear memory. Hippocampus, 15(7), 874–880.

Poppenk, J., Evensmoen, H. R., Moscovitch, M., & Nadel, L. (2013). Long-axis specialization of the human hippocampus. Trends in Cognitive Science, 17(5), 230–240.

Qasim, S. E., Miller, J. F., Inman, C., Gross, R. E., Willie, J., Bradley, L., … Joshua, J. (2019). Memory retrieval modulates spatial tuning of single neurons in the human entorhinal cortex. Nature Neuroscience, 1–9.

Raghavachari, S., Kahana, M. J., Rizzuto, D. S., Caplan, J. B., Kirschen, M. P., Bourgeois, B., … Lisman, J. E. (2001). Gating of human theta oscillations by a working memory task. Journal of Neuroscience, 21(9), 3175–3183.

Ramos-Loyo, J., & Sanchez-Loyo, L. (2011). Gender differences in eeg coherent activity before and after training navigation skills in virtual environments. Human Physiology, 37(6), 700–707.

Rivas, J., Gaztelu, J. M., & García-Austt, E. (1996). Changes in hippocampal cell discharge patterns and theta rhythm spectral properties as a function of walking velocity in the guinea pig. Experimental Brain Research, 108(1), 113–8.

Royer, S., Sirota, A., Patel, J., & Buzsaki, G. (2010). Distinct representations and theta dynamics in dorsal and ventral hippocampus. The Journal of neuroscience, 30(5), 1777–1787.

Schmidt, B., Hinman, J. R., Jacobson, T. K., Szkudlarek, E., Argraves, M., Escabí, M. A., & Markus, E. J. (2013). Dissociation between dorsal and ventral hippocampal theta oscillations during decision-making. Journal of Neuroscience, 33(14), 6212–6224.

Schultheiss, N. W., Schlecht, M., Jayachandaran, M., Brooks, D. R., McGlothan, J. L., Guilarte, T. R., & Allen, T. A. (2019). ‘awake delta’ and theta-rhythmic hippocampal network modes during intermittent locomotor behaviors in the rat. bioRxiv.

Seidenbecher, T., Laxmi, T., Stork, O., & Pape, H. (2003). Amygdalar and hippocampal theta rhythm synchronization during fear memory retrieval. Science, 301(5634), 846.

Siapas, A. G., Lubenov, E. V., & Wilson, M. A. (2005). Prefrontal phase locking to hippocampal theta oscillations. Neuron, 46(1), 141–151.

Sirota, A., Montgomery, S., Fujisawa, S., Isomura, Y., Zugaro, M., & Buzsaki, G. (2008). Entrainment of Neocortical Neurons and Gamma Oscillations by the Hippocampal Theta Rhythm. Neuron, 60(4), 683–697.

Sławińska, U., & Kasicki, S. (1998). The frequency of rat’s hippocampal theta rhythm is related to the speed of locomotion. Brain Research, 796(1-2), 327–331.

Staudigl, T., & Hanslmayr, S. (2013). Theta oscillations at encoding mediate the context-dependent nature of human episodic memory. Current Biology, 23(12), 1101–1106.

Strange, B. A., Witter, M. P., Lein, E. S., & Moser, E. I. (2014). Functional organization of the hippocampal longitudinal axis. Nature Reviews Neuroscience, 15(10), 655–669.

Vass, L. K., Copara, M. S., Seyal, M., Shahlaie, K., Farias, S. T., Shen, P. Y., & Ekstrom, A. D. (2016). Oscillations go the distance: Low-frequency human hippocampal oscillations code spatial distance in the absence of sensory cues during teleportation. Neuron, 89(6), 1180–1186.

von Stein, A., Rappelsberger, P., Sarnthein, J., & Petsche, H. (1999). Synchronization between temporal and parietal cortex during multimodal object processing in man. Cerebral Cortex, 9(2), 137–150.

Voytek, B., Canolty, R., Shestyuk, A., Crone, N., Parvizi, J., & Knight, R. (2010). Shifts in gamma phase–amplitude coupling frequency from theta to alpha over posterior cortex during visual tasks. Frontiers in Human Neuroscience, 4.

Watrous, A. J., Fried, I., & Ekstrom, A. D. (2011). Behavioral correlates of human hippocampal delta and theta oscillations during navigation. Journal of Neurophysiology, 105(4), 1747–1755.

Watrous, A. J., Lee, D. J., Izadi, A., Gurkoff, G. G., Shahlaie, K., & Ekstrom, A. D. (2013). A comparative study of human and rat hippocampal low-frequency oscillations during spatial navigation. Hippocampus, 23(8), 656–661.

Watrous, A. J., Miller, J., Qasim, S. E., Fried, I., & Jacobs, J. (2018). Phase-tuned neuronal firing encodes human contextual representations for navigational goals. eLife, 7, e32554.

Watrous, A. J., Tandon, N., Conner, C. R., Pieters, T., & Ekstrom, A. D. (2013). Frequency-specific network connectivity increases underlie accurate spatiotemporal memory retrieval. Nature Neuroscience, 16(3), 349–356.

Winson, J. (1978). Loss of hippocampal theta rhythms in spatial memory deficit in the rat. Science, 201, 160–163.

Wyble, B. P., Hyman, J. M., Rossi, C. A., & Hasselmo, M. E. (2004). Analysis of theta power in hippocampal eeg during bar pressing and running behavior in rats during distinct behavioral contexts. Hippocampus, 14(5), 662–674.

Yassa, M. A. (2018). Brain rhythms: higher-frequency theta oscillations make sense in moving humans. Current Biology, 28(2), R70–R72.

Zhang, H., & Jacobs, J. (2015). Traveling theta waves in the human hippocampus. The Journal of Neuroscience, 35(36), 12477–12487.

Zhang, H., Watrous, A. J., Patel, A., & Jacobs, J. (2018). Theta and alpha oscillations are traveling waves in the human neocortex. Neuron, 98(6), 1269–1281.e4.

